# DNA copy number imbalances in primary cutaneous lymphomas (PCL)

**DOI:** 10.1101/417766

**Authors:** Georgiana Gug, Qingyao Huang, Elena Chiticariu, Caius Solovan, Michael Baudis

## Abstract

Cutaneous lymphomas (CL) represent a clinically defined group of extranodal non-Hodgkin lymphomas harboring heterogeneous and incompletely delineated molecular aberrations. Over the past decades, molecular studies have identified several chromosomal aberrations, but the interpretation of individual genomic studies can be challenging.

We conducted a meta-analysis to delineate genomic alterations for different types of PCL. Searches of PubMed and ISI Web of Knowledge for the years 1996 to 2016 identified 32 publications reporting the investigation of PCL for genome-wide copy number alterations, by means of comparative genomic hybridization techniques and whole genome and exome sequencing. For 449 samples from 22 publications, copy number variation data was accessible for sample based meta-analysis. Summary profiles for genomic imbalances, generated from case-specific data, identified complex genomic imbalances, which could discriminate between different subtypes of CL and promise a more accurate classification. The collected data presented in this study are publicly available through the “Progenetix” online repository.

## Introduction

Cutaneous lymphomas (CL) represent a complex group of extranodal non-Hodgkin lymphomas for which the underlying molecular events are incompletely understood. So far, the classification of cutaneous lymphoma remains one of the most challenging areas of oncologic dermato-pathology. The first widely accepted classification, which emphasizes the features of cutaneous lymphomas as primary tumors of the skin, was the “WHO / EORTC classification (2005)”, representing a consensus based on the European Organization for Research and Treatment of Cancer (EORTC) and World Health Organization (WHO) classifications (1) (2). All forms of primary CL found in the WHO / EORTC consensus have been adopted by the WHO classifications (3). Updates which integrate these nosological entities into a larger classification, containing both nodal and extranodal lymphomas as different categories, are made periodical, with currently active classification having been published in 2016 (4). Since our metaanalysis is based on data spanning the period from 1996 to 2016, we decided to follow the 2008 classification to avoid discrepancies and ambiguities in code assignment.

CLs are represented by primary cutaneous T cell lymphomas (CTCLs) in approximately 65% of cases and in a smaller percentage by primary cutaneous B cell lymphomas (CBCL). Although a large number of clinicopathologic variants are being acknowledged in the WHO classification, a clear, biologically supported definition of specific entities is missing so far.

CTCL identifies a group of extranodal T-cell lymphomas characterized by the infiltration of malignant CD4+ T-cells in the skin (5). Among those, mycosis fungoides (MF) represents the most common diagnosis (44% of CL) (6). MFs are neoplasms of skin-homing T cells with CD4+ T-helper phenotype and Th2 pattern which commonly behave as a lowgrade lymphoma with a protracted clinical evolution; however, a subset of patients progress rapidly to severe forms, refractory to current treatment modalities (7). For MF, a wide spectrum of subtypes has been described, such as folliculotropic, pagetoid reticulosis and granulomatous slack skin (8). In contrast to the frequently indolent MF, Sézary syndrome (SS) represents a rare aggressive subtype of CTCL defined by diffuse pruritic rash, lymphadenopathy and the early appearance of malignant T-cells in the peripheral blood. The mechanisms underlying the proliferation of neoplastic CD4+ T-cells in SS are not fully understood.

Lymphomatoid papulosis and primary cutaneous anaplastic large cell lymphoma are part of a spectrum of CD30+ cutaneous lymphoproliferative diseases, the second most common group of CTCL. They have similar morphologic and immunophenotypic characteristics, with differentiation relying predominantly on the clinical representation. Adult Tcell leukaemia / lymphoma is an aggressive peripheral Tlymphocytic neoplasm caused by a human retrovirus, human T-cell lymphotropic virus type 1 (HTLV-1). Skin involvement is generally a manifestation of disseminated disease, but a slowly progressive form which may have primary cutaneous involvement has been described (smoldering subtype) (9).

Other recognized CTCLs include subcutaneous panniculitis-like T-cell lymphoma, extranodal NK / T-cell lymphoma, and peripheral T-cell lymphomas, rare subtypes (Cutaneous gamma/delta T Cell Lymphoma, Primary cutaneous CD4+ small-medium sized pleomorphic T-Cell lymphoma, Primary cutaneous aggressive epidermotropic CD8+) (3).

The class of cutaneous B cell lymphomas (CBCL) have provided cause for debate for a long time, and present a compelling reason for developing a new classification. Currently recognized categories include Primary Cutaneous Marginal Zone B-cell lymphoma, Primary Cutaneous Follicle Centre Lymphoma, primary cutaneous diffuse large B-cell lymphoma, and primary cutaneous intravascular large B-cell lymphoma (3).

To date, little is known about the underlying molecular events in CL, both because of the limited ability to extrapolate lessons learnt from the nodal counterpart (similar pathological entities with different biological behavior, prognosis and treatment) and also the difficulty to conduct studies addressing CL (overall low incidence together with high clinicopathologic heterogeneity) (10). Over the past decade, molecular studies have identified several molecular events such as gene specific mutations, aberrant gene expression profiles, upregulated micro-RNAs, telomere shortening, and chromosomal aberrations (11–18). However, the etiology remains unknown for the majority of cases, with a set of common pathways or initiating events in different stages of lymphocyte maturation to be identified. Importantly, the behavior of a malignant clone of lymphocytes is not only the result of its uncontrolled proliferation but also may depend on factors and stimulants from the cutaneous microenviroment.

A number of studies have been investigating structural genome variants in CL, such as chromosomal copy number alterations, localized copy number variations, deletions, amplifications and insertions. While these genome variants may be based on different rearrangement mechanisms, the common end point of somatic copy number variations/alterations (CNV) is a potential gene dosage effect, promoting over- or underrepresentation of the transcripts and proteins derived from genes in the affected regions. While large CNVs make it difficult to identify the possible target gene(s), “focal CNV” an operational term based on a limited size up to 3Mb and the inferred relevance of direct gene targeting have the potential to point directly towards pathogenetic gene defects (19).

The increasing number of genomic studies and the advances in genome analysis technologies promise a better understanding of the molecular events responsible for clonal transformation and histopathological representation of CL. Also, genomic alterations have the potential to support a better classification of these diseases and can be supportive in prognostic evaluation and treatment response. However, the interpretation of individual genomic studies especially in rare diseases can be challenging due to limitations such as small sample sizes as well as differences in analysis methods and data processing. Here, we conduct a meta-analysis of published CL studies to identify genomic alterations and alteration patterns specific for different types of CL. The collected data presented in this study are publicly available through the “Progenetix” repository for molecular-cytogenetic data.

## Methods

### Search strategy

We conducted a search of published research using PubMed and ISI Web of Knowledge from the year 1996 to 2016 with no language restrictions. Search terms included the following sets of keywords variably combined: “lymphoma”, “cutaneous lymphoma”, “skin lymphoma”, “CTCL”, “mycosis fungoides”, “sezary”, “leukaemia”, “leukemia”, “lymphomatoid papulosis”, and “cgh” or “CGH” or “aCGH” or “comparative genomic hybridisation” or “snp” or “SNP” or “array” or “genomic array” or “genome” or “copy number” or “dna microarray” or “amplification”. Additional, we followed up on references from the selected articles for possibly relevant publications followed by evaluation of the abstracts. If more than one article was published using the same series of subjects, we chose the latest or the most complete study for this meta-analysis.

### Inclusion and exclusion criteria

For selection of the studies we followed the guidelines of the critical checklist proposed by the Dutch Cochrane Centre Meta-analysis of Observational Studies in Epidemiology (MOOSE) (20). Articles were identified as eligible when they fit the following criteria: i. they contain genomic copy number data with whole genome coverage and access to this information; ii. diagnosis of Primary Cutaneous Lymphoma (PCL) with equivalent terminology in EORTC-WHO (2005) and WHO (2008) classifications (Table 1); iii. matching available or convincingly inferred locus information.

**Table 1.**
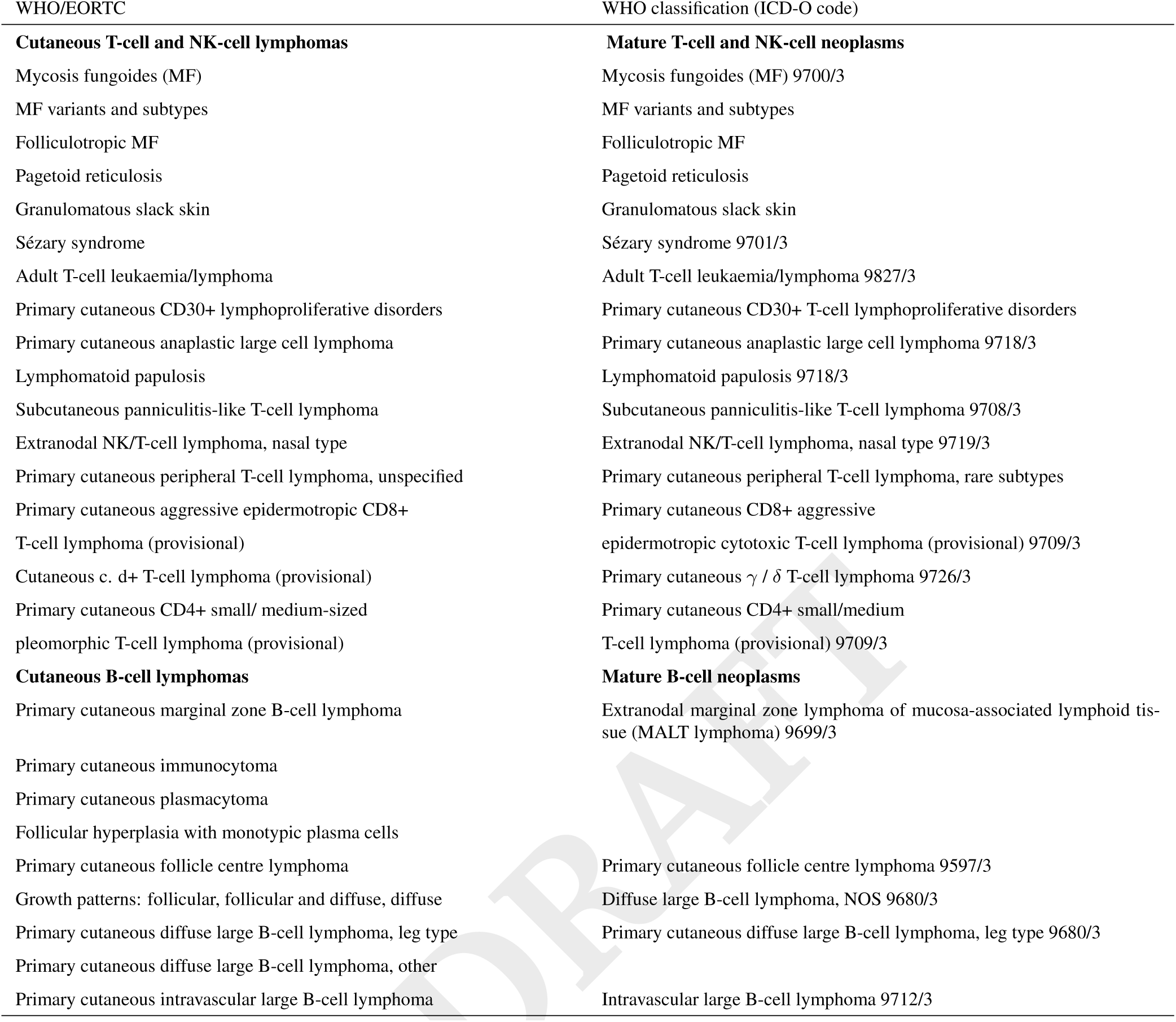
WHO/EORTC (2005) and WHO (4th edn., 2008) classifications of CL

The process of study retrieval is presented in Table 2 according to PRISMA (preferred reporting items for systematic reviews and meta-analyses) statement.

**Table 2.** Study retrieval

| All | Full text | Open access | After excluding the doubles | After article review | After data review |
| --- | --- | --- | --- | --- | --- |
| 138752 | 120354 | 46754 | 23266 | 32 | 22 |

### Data extraction and quality assessment

For each of the identified studies, the following information was extracted: first author, year of publication, population demographics, clinical characteristics, follow-up data where available, and genomic alterations for each patient individually. The techniques considered for the assessment of genomic copy number aberrations included both chromosomal and array-based comparative genomic hybridisation (cCGH/aCGH).

Copy number aberration data was either mapped to the genomic base locations of a 862 band karyotype data (UCSC genome browser hg38 mapping; genome.ucsc.edu) or based on the genome mapped segmentation calls where available. For genomic array data without called regional gain/loss information, we used the arrayMap software tools to segment and visualize the probe specific data and derived regions of imbalances.

### Data analysis

We used the complete DNA copy number alterations (CNAs) data to generate non-overlapping genomic segments, in order to evaluate overall and subgroup specific frequency as well as sample based patterns of regional copy number changes. Case-specific CNA data was visualized and ordered by using data matrices containing imbalance status (gain/loss) mapped to 1Mb genomic intervals. Hierarchical clustering of matrices was used to order the cases, the obtained order then used for re-plotting of CNA in their original resolution. The relative genomic CNA coverage (number of bases with CNA compared to genome size) was used as a proxy for CNA complexity. The imbalance distribution was determined, by calculating the gain/loss frequencies, mapped to genomic intervals on a 1Mb level. A heatmap of gain/loss distributions was generated to compare the copy number profiles between samples. For patients with clinical follow-up, a limited survival analysis was performed.

## Results

Our search using the keywords with a logical “OR” assertion returned a total of 138752 abstracts. Title and abstract review resulted in the exclusion of 115486 papers which were duplicates and another 23234 which were not relevant to our study. Additionally, no full text was available for 18398 publications, which were thus excluded. We reviewed 32 articles in full. Ten articles were excluded due to the lack of individual data (even after contacting the authors) or because they didn’t meet the inclusion criteria (secondary cutaneous etc.). Finally, we included 22 articles in the present meta-analysis. The 22 studies included here, comprised 449 samples in total (Table 3).

**Table 3.** Cutaneous lymphoma samples included in meta-analysis.

| WHO classification | Diagnostic text | ICD-O | No. Samples | PMID |
| --- | --- | --- | --- | --- |
| Mycosis fungoides (MF) | Mycosis fungoides | 9700/3 | 145 (32.2%) | 18558289, 18832135, 26551667, 17584911, 15086538, 18663754, 12207585, 19759554 |
| Sézary syndrome (SS) | Sézary syndrome | 9701/3 | 142 (31.6%) | 10084322, 10874879, 12207585, 12557225, 18413736, 18558289, 19843862, 25314094, 26551667, 15086538 |
| Primary cutaneous anaplastic large cell lymphoma (ALCL) | Anaplastic large cell lymphoma [cutaneous];<br>Anaplastic large cell lymphoma, cutaneous type | 9718/3 | 73 (16.2%) | 12648233, 12696066, 14595754, 15086538, 15111330, 19710685, 26551667 |
| Primary cutaneous peripheral T-cell lymphoma, rare subtypes * (TCRS) | Peripheral T-cell lymphoma [cutaneous] (PTL-NOS) | 9709/3 | 16 (3.5%) | 15111330, 19710685, 26551667 |
| Extranodal marginal zone lymphoma of mucosa-associated lymphoid tissue (MALT lymphoma) (MZBL) | Extranodal marginal zone B-cell lymphoma;<br>Marginal zone lymphoma [cutaneous] | 9699/3 | 16 (3.5%) | 12203778 |
| Primary cutaneous follicle centre lymphoma (FCBL) | Follicular lymphoma [primary cutaneous];<br>Follicular lymphoma [cutaneous] | 9597/3 | 19 (4.2%) | 12203778, 15175042, 16274455 |
| Diffuse large B-cell lymphoma ‡(DLBCL) | Diffuse Large B-cell Lymphoma [extranodal, skin];<br>Diffuse large B-cell lymphoma [primary cutaneous, leg type];<br>Diffuse large B-cell lymphoma [cutaneous]; | 9680/3 | 38 (8.4%) | 16274455, 15175042, 12203778 |

\* Cutaneous peripheral T-cell lymphoma, unspecified in EORTC/WHO classification; 9709/3 defines three subtypes: Primary cutaneous CD4-positive small/medium T-cell lymphoma (CD4+ SMTL), Primary cutaneous CD8-positive aggressive epidermotropic T-cell lymphoma (CD8+ AECTCL), and Primary cutaneous gamma/delta T-cell lymphoma (CGD-TCL).
Primary cutaneous marginal zone B-cell lymphoma in EORTC/WHO classification ‡9680/3 defines two subtypes in WHO classification: Diffuse large B-cell lymphoma, NOS and Primary cutaneous diffuse large B-cell lymphoma, leg type; Primary cutaneous diffuse large B-cell lymphoma, leg type and Primary cutaneous diffuse large B-cell lymphoma, other in EORTC/WHO classification.

### A. Distribution of studies

Almost all of the studies included in our meta-analysis originated in Europe; one case was selected from a Chinese paper and one additional study was the result of a collaboration between groups from the USA and the Netherlands (41 patients with T-cell lymphoma). No studies from other geographic regions could be included as they didn’t met the inclusion criteria.

### B. CNAs in CL

The genomic abnormalities found in CTCL (376 samples) and CBCL (73 samples) are summarized in Figure 1.

**Fig. 1.**
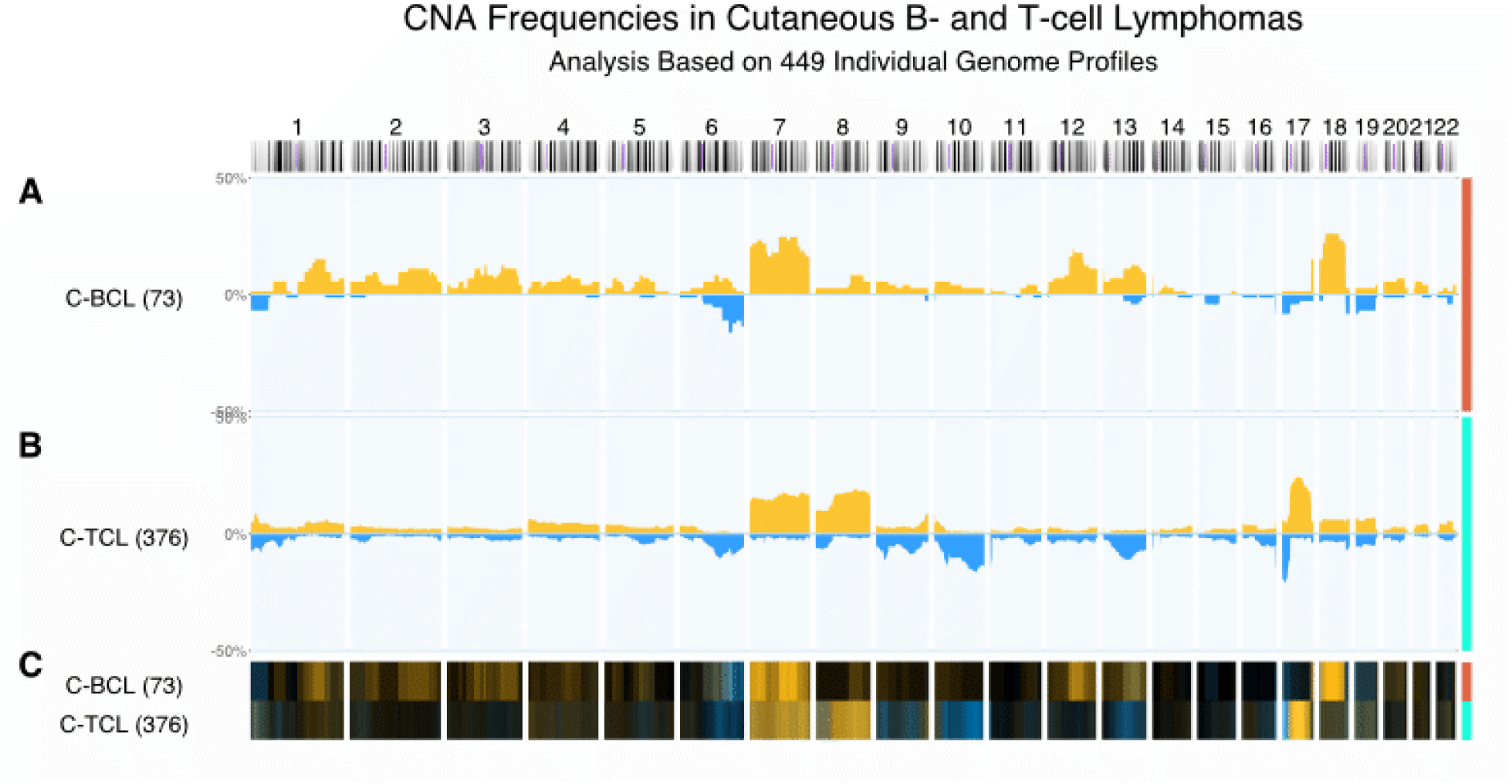
Frequency of genomic gains and losses in C-BCL and C-TCL. This histogram shows the frequency of genomic gains/amplifications (orange, up) and deletions (blue, down) along genomes of cutaneous B(A) and T-CL (B), ordered from chromosome 1 to 22. In this plot, a frequency of e.g. 25% for a gain means that in 25% of the selected cases, a genome duplication was observed in 25% of the samples, for the matching genomic interval. The same situation is also valid for deletions. In panel **C**, the regional changes are compared using a heatmap style display, in which a high local frequency of copy number gains and losses results in predominantly yellow and blue color, respectively.

Reviewing the studies on CTCL included in our metaanalysis, we found that the most frequent CNAs, i.e. genomic regions with frequent imbalances, consisted of copy number gains on chromosome 17q, 7 and 8q, while the most frequent losses presented themselves in regions on 17p, 6q, 9p21 and 13q. Additionally, recurring deletions could be observed on 1p, 8p, and chromosome 10q. Not surprisingly, each of these regions has been found to harbour oncogenes or tumor suppressor genes, respectively, with critical roles in the course and development of these oncologic entities. Karenko *et al.* evaluated the clinical evolution of CTCL for an average 54 months in comparison with chromosomal abnormalities. They analyzed chromosomes 1, 6, 8, 9, 11, 13/21, 15 or 17 from 5 cases of Large Plaque Parapsoriasis (LPP), 8 MF and 2 SS, using G-banding and in situ hybridization techniques. Chromosomal aberrations from patients achieving complete clinical remission predominantly involved chromosomes 1, 6 and 11, while patients with active and progressing disease showed mutation on chromosomes 1, 6, 8, 11 and 17 (21).

In the histogram of CBCL gains are harbouring loci like 12q, 18q and losses in 6qter. While in the genomic copy number gains represented the more prevalent type of CNA in both Band T-cell derived cutaneous lymphomas, the involved genomic regions differed in accordance with the lineage. The most frequent gains in T-NHL involved 17q and 8q, while the most frequent duplications in B-NHL occurred on 18q (involving the BCL2 and MALT1 locus) and as frequently focal amplification on 12q21 (around CDK4, oncogenes GLI1 and MDM2).

### C. CNAs in MF

Mycosis fungoides, a malignancy of skin homing T cells, is the most common type of CTCL whose early diagnosis is often challenging. The use of genomic analyses has been studied and its importance is undeniable. Gains in copy numbers were more often described than deletions in this type of CL. Recurrent alterations were highlighted as gains on chromosome 1, 7, 8, 9qter, 17, 19, 22 and losses of 6q, 9pter, 13q and 17p (22).

Figure 2A, generated from 145 samples of MF, highlights duplications on chromosome 1p, chromosome 7, 8 with (predominantly 8qter), 9qter, 17 predominant on the long arm, 19, 22; and deletions on 6q, 10, 13q, 16q, 19, 9p and q (2 predominant regions, with narrow hot spot on 9p21.3) and 17p. The TP53 gene, also known as the guardian of the genome, is an important instruction for p53 protein, which acts as a tumor suppressor. This gene is located on 17p13.1 and its deletion is easily noticed on MF histoplot and also on the histoplots generated for CTCL and CBCL (Figure 1). The deletion spike seen in 9p21.3 is correspondent to CDKN2A, CDKN2B and MTAP genes, that encode tumor suppressor proteins p14 (role in p53 protection) and p16. Gains seen in the long arm of chromosome 8 on locus 8q24.21 are harboring the MYC proto-oncogene, which is an functional target in numerous human cancers (23).

**Fig. 2.**
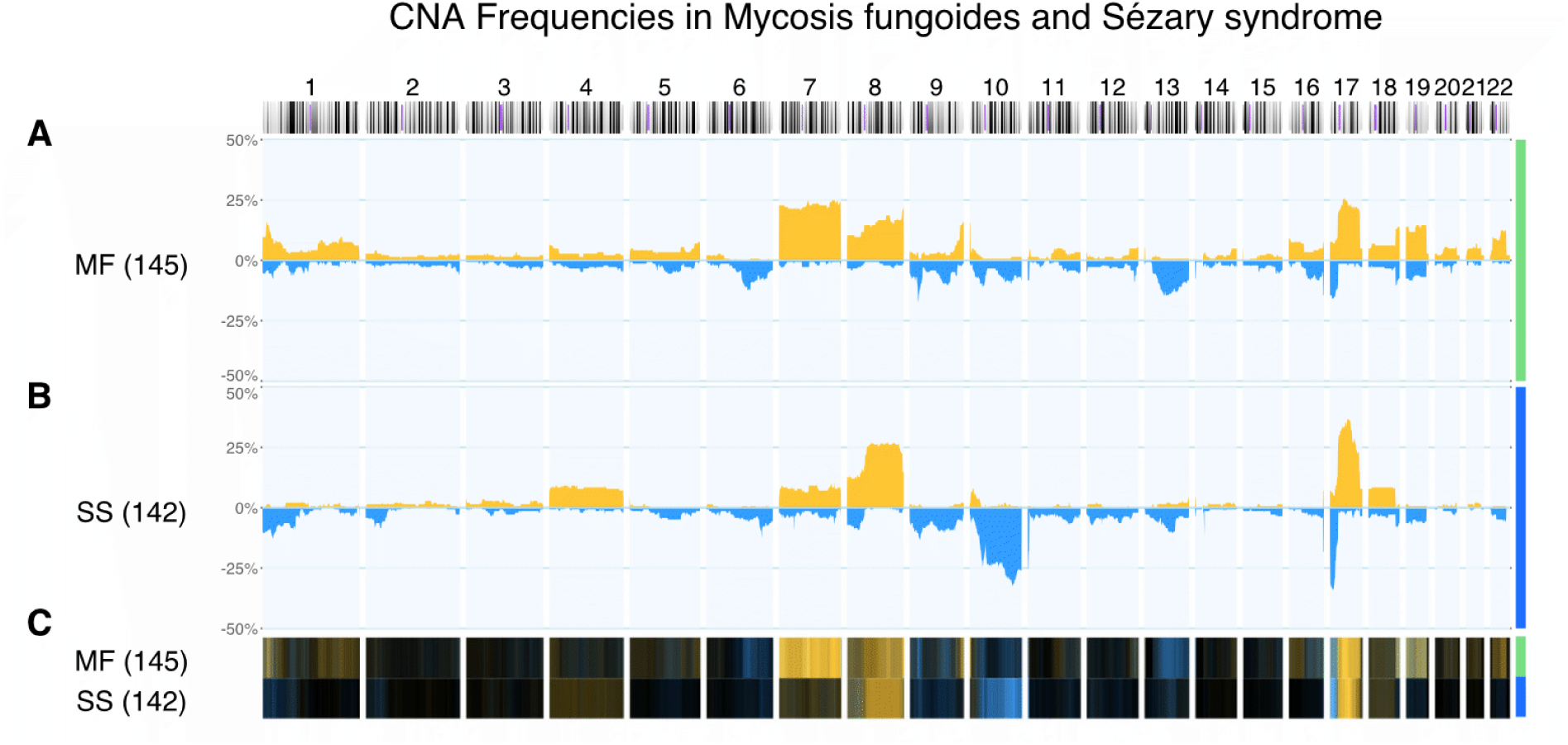
Comparison of CNAs in MF and SS. In **A**, the distribution of local CNAs in MF is shown, and compared to those in SS in **B**.

### D. ALK amplification in a case of MF

Reviewing the data from the van Doorn *et al.* paper, we observed in one case of MF (GSM325151) a gain of the distal part of chromosome 2p, with a breakpoint in or close to the 3’ end of the ALK proto-oncogene. Also, while the probe segmentation resulted in a single segment (2:29193223-29920658 DUP; hg38 coordinates), a cluster of 5 probes covering the ALK locus suggested for an additional copy number gain. While typically ALK is implicated in the pathogenesis of anaplastic large cell lymphoma (ALCL; see SubsectionG) (24, 25) or Diffuse large B-cell lymphoma (DLBCL; see Subsection K) (26), this observation provides a rare report of ALK rearrangement in MF (Figure 3).

**Fig. 3.**
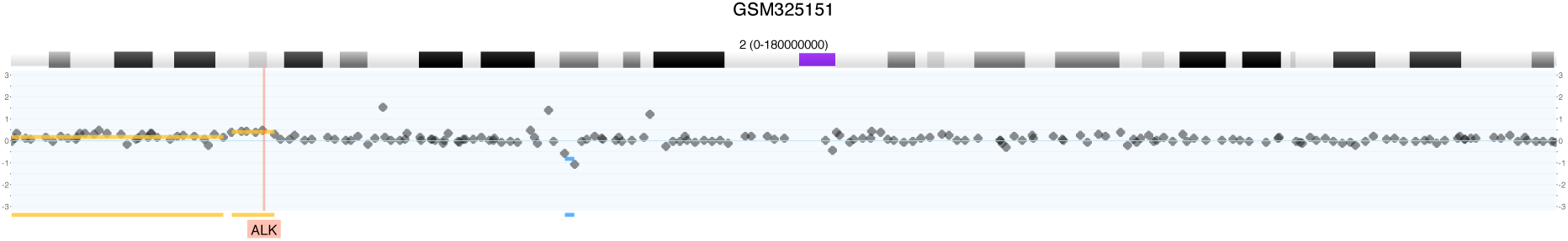
ALK rearrangement and suspected amplification in a case of MF.

In histological analysis, transformed MF presents with similarities to primary cutaneous ALCL, although ALK expression does not constitute a typical feature of MF. After Covington *et al.* identified a case of MF and concurrent nodal ALCL with the expression of ALK (27), they decided to analyze 103 biopsies of 96 MF patients for ALK expression. Out of all biopsies, only the case with MF and nodal ALCL association presented positive for ALK, leading to the conclusion that ALK is not routinely present in any stage or variant of MF. In another report one case of CTCL with an aggressive evolution turned ALK positive (28). The authors concluded that most likely this case represented CD30+ transformed MF. There, ALK expression was accompanied by an aggressive course in contrast to the typical favorable prognosis of ALK+ ALCL. The rarity of such observations emphasizes the value of data accessibility for meta-analyses, thereby allowing the integration and proper evaluation of such rare genomic events, since multiple instances of single observations may support recurring pathogenetic mechanisms.

### E. CNAs in SS

Sézary Syndrome, although clinically related to MF and sometimes genetic and molecular aberrations are described with similar patterns (29), it has a negative impact on the survival and shares distinct genetic properties. In Figure 2B, generated from 142 samples of SS, gains on chromosome 4, 7, 19 and predominant on chromosome 8q and 17q and losses on 1p, 2p, 6q, 8p, 9, 13q with massive losses on chromosome 10 and a spike on 17p (TP53 locus).

### F. MF versus SS CNAs

We compared CNAs between MF and SS, to spot the differences and compare the data with literature reports. Figure 2C compares the distributions of CNAs in a nearly even set of total 287 samples, collected from 14 publications.

Main differences between MF and SS in our 287 samples are seen on chromosomes:

• 1p (MF gains, SS losses)

• 2 (losses on short armSS)

• 4 (gains in some SS),

• 6q (losses predominant in MF),

• 7 (much higher number of gains in MF),

• 9qter (gains in MF),

• 10 (very high no. of losses in SS; overall a rare event in cancer apart from gliomas),

• 13 (higher no. of deletions in MF),

• 17 (higher no. of gains in q arm and losses in p arm in SS, probably reflecting i17q and TP53 involvement),

• 19 (gains in MF)

The loss in TP53 gene located on 17p13.1 is more prominent in SS than MF. The deletion spike seen in 9p21.3 corresponding to the locus of the CDKN2A, CDKN2B and MTAP genes is encountered in both MF and SS with a slightly higher rate in MF. Gains seen in 8q24.21, encoding the MYC proto-oncogene, are somewhat more frequent in SS.

Sézary Syndrome is characterized by highly recurrent alterations including gains on 17q23-25 and 8q24 as well as losses on 17p13, with frequencies higher than those in MF (30). In our CNA analysis of this loci, both SS and MF are affected by these CNA, but with higher numbers in SS. In the same paper it was reported that gains in 7q36 rarely occur in SS, in contrast to MF. In our analysis, gains of mostly the whole chromosome 7 appear in both diseases but predominantly MF. Overall, in accordance with van Doorn *et al.* (30):

• MF characterized by gains on chromosome 1 and 7 and loses on chr. 9

• SS characterized by gains on chromosome 8 and 17 and loses on chr. 10

The differences in focal gains and losses between MF and SS highlight the unique character of these pathologies, substantiating a differentiated approach of their classification and clinical evaluation.

### G. CNAs in ALCL

Cutaneous ALCL has an usual indolent clinical evolution and overall good prognosis. In our study, cALCL was characterized by gains on chromosome 1, 2p, 5, 6p, 9 and high no. of gains on chromosome 7. Frequent deletions are seen in chromosome 6 (mostly on the long arm), 13 but also some deletions on 1p, 3p, 16q, 17p, 18p. All can be observed on the graphic of 73 cALCL cases (Figure 4B). Oncogenes like NRAS (1p13.2), RAF1 (3p25), CBFA2 (21q22.3), JUNB (19p13.2) are amplified more or less in the 73 samples of ALCL. Even if ALK gene rearrangements are rarely seen in cALCL, a subset of case reports describes this gene rearrangement (24). In our case, copy number gains are positive on the ALK locus gene 2p23.2-p23.1 (in 4 cases out of 73). Deletions in TP53 gene located on 17p13.1 and PRDM1 (6q21) is also seen in our cases, fact reported by other authors as often events in this type of lymphoma (25). This consistent copy number alteration is an important pawn for future studies in those cases with severe clinical evolution.

**Fig. 4.**
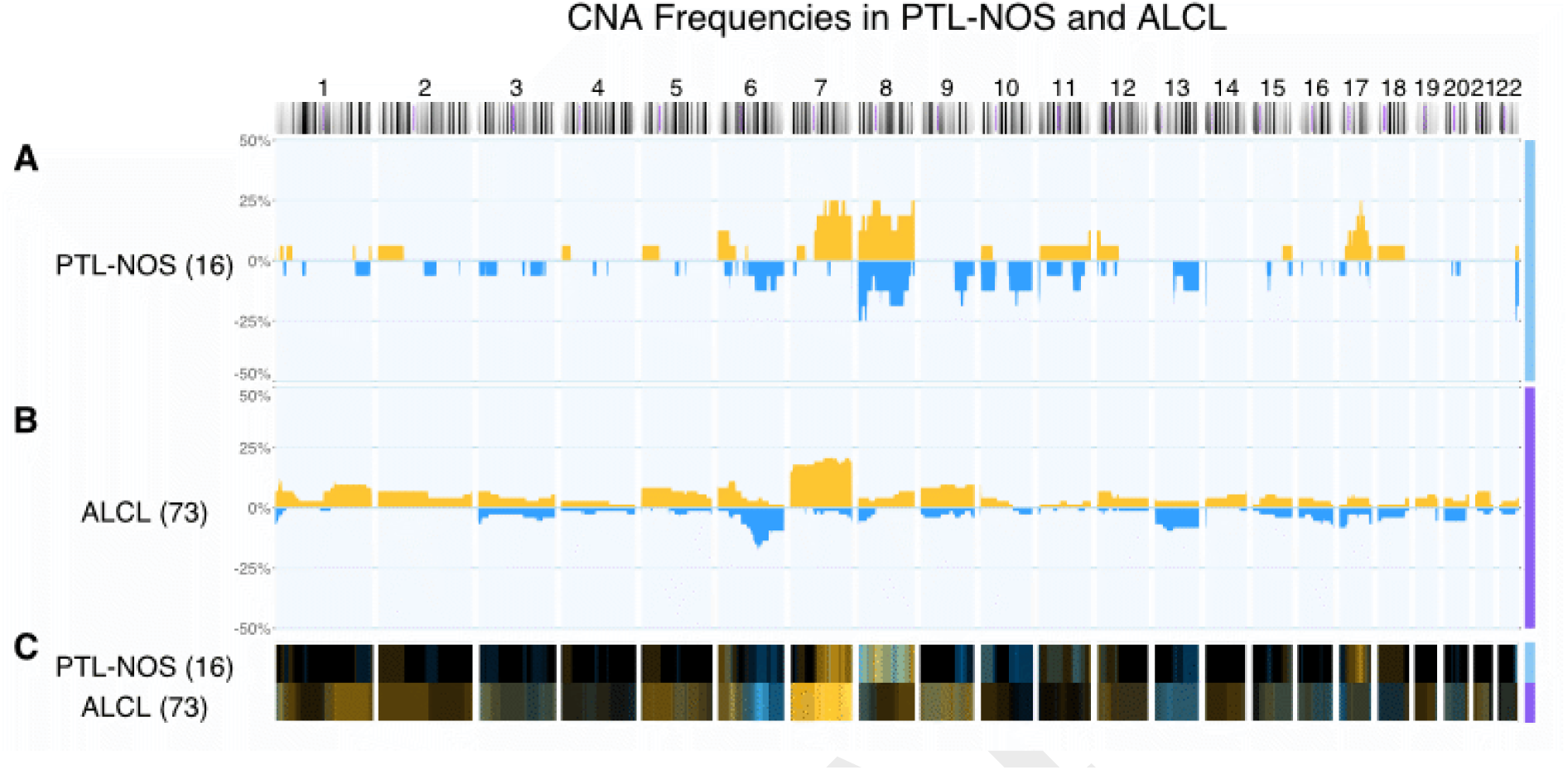
CNAs in PTL-NOS and ALCL.

### H. CNAs in TCRS (PTL-NOS)

In contrast with ALCL, primary cutaneous peripheral T-cell lymphoma not otherwise specified (PTL-NOS) shows an aggressive clinical evolution and bad prognosis. PTL-NOS shows gains on 6p, 7q and 17q, and was distinguished by high no. of gains on chromosome 8. Losses are encountered on 6q, 9q, chromosomes 10, 11, 13 and also some deletions on various regions of chromosome 8 (Figure 4A). However, this characterisation may be influenced by the low number of samples (16 cases).

### I. CNAs in MALT lymphoma (MZBL)

Out of the 16 samples of MALT lymphoma (MZBL) accessible, 6 presented CNAs (only in gains, recurrently on chromosome 7, 8q, 13q, 18p, 20p, 21; Figure 5C). Gains on 1p22.3 corresponding to BCL10 apoptosis gene is present in 2 cases. This is reported as being associated with the development of extracutaneous disease but without importance in primary cutaneous disease (31).

**Fig. 5.**
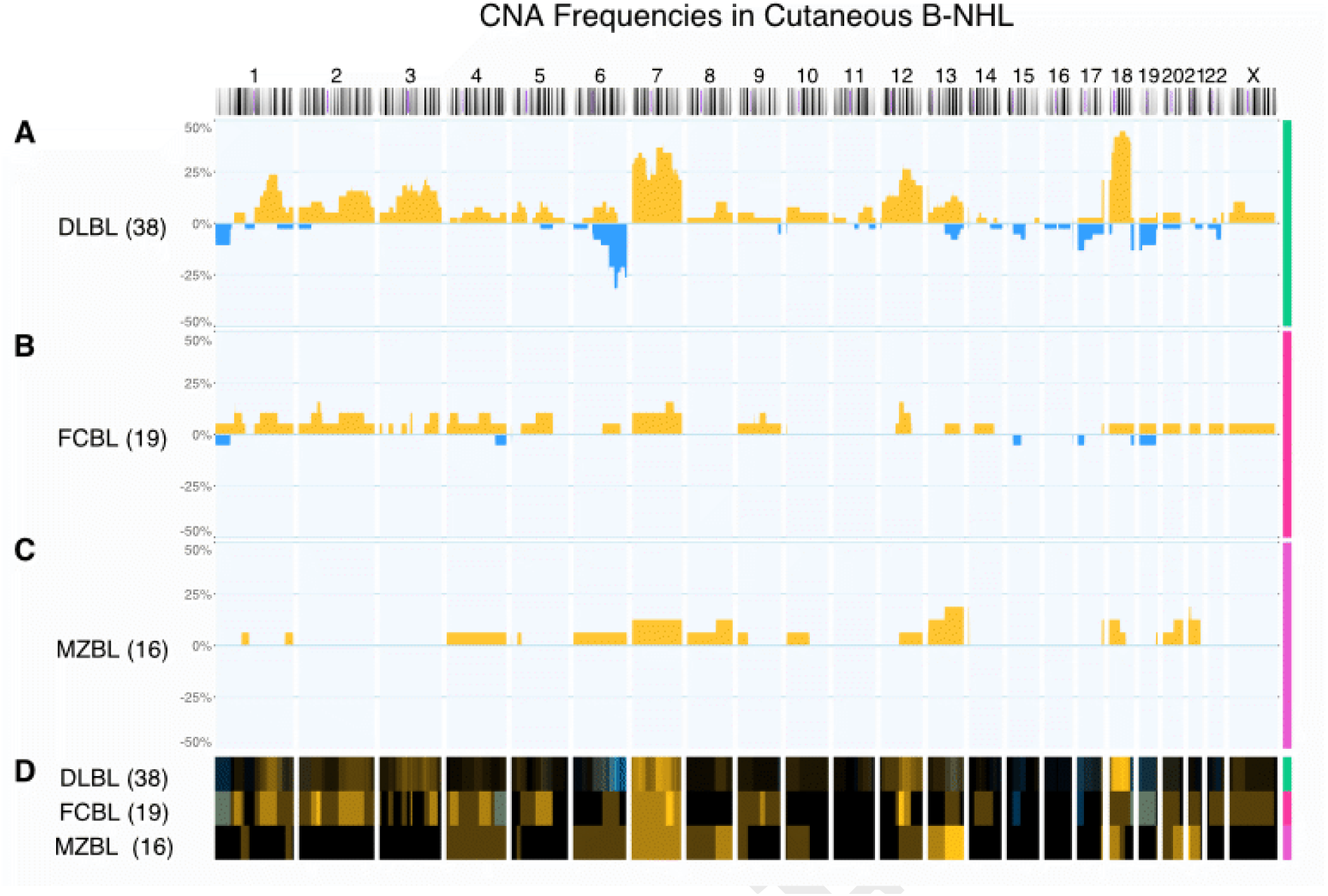
CNAs in cutaneous B-NHL.

### J. CNAs in FCBL

Out of 19 samples of FCBL, 6 presented CNAs mostly represented by gains. Most often gain was on chromosome 1p22, 1q21-1q31, 2p16 with a spike on 2p13, 2q21-q32, 3q25-qter, chromosome 4, chromosome 5, chromosome 7 with high number of gains in 7q31-q32, chromosome 8, 12(Figure 5B). Literature reports reveal amplification in 2p16.1 in most cases of FCL with amplification of c-REL apoptotic gene, case confirmed by our analysis. Deletions of 14q31.33 were also highlighted, situation not met in the current meta-analysis (32).

### K. CNAs in DLBCL

From all CBCL, DLBCL has the worst prognosis and survival. Genetic studies revealed recurrent amplifications in 18q21.31q21.33, which includes the MALT1 and BCL2 loci, as well as increased expression of proto-oncogenes PIM1(6p21.2), WDR26 (1q42.11-q42.12), MYC(8q24.21) and IRF4(6p25.3). Recurrent deletions were described in 9p21.3 (CDKN2A, CDKN2B, NSG-x genes) (32).

In Figure 5A, in 38 DLBCL case analysis, we observe a large variety of copy number alteration in this type of lymphoma. CNAs are mostly covered by gains. They are encountered on chromosome 1q, 2, 3, 6, 8q and 13 and a large number of amplifications in chromosome 7, 12 and 18. Less common losses are observed on 1p, 6q with a spike on 6q22 and on chromosomes 17 and 19 with higher frequency on short arms. Wiesner *et al.* reported the most frequent numerical aberrations in their 40 cases analysis. From the most frequent to the less frequent, gains affected chromosome 12, 7, 3, 18q, 11 and losses affected chromosome 18. In our cohort of 38 samples, most frequent gains were on chromosome 18q (including BCL2 gene and MALT1 gene locus), 7q, 7p, 1q (including WDR26 gene amplifications), 12q, 3q, 2q, 13q, 6q, 8q (including MYC gene amplification). Most frequent losses were present on chromosome 6q, 17p, 19p, 20q, 6p, 1p. Deletions on 9p21 locus corresponding to CDKN2A gene were not met in our patient group.

### L. Genomic imbalance patterns

To quantitatively measure genomic imbalances, we derived the fractions of genomic regions with aberrant copy number per sample (abfrac) and determined their distributions among all the analyzed CL entities (Figure 6).

**Fig. 6.**
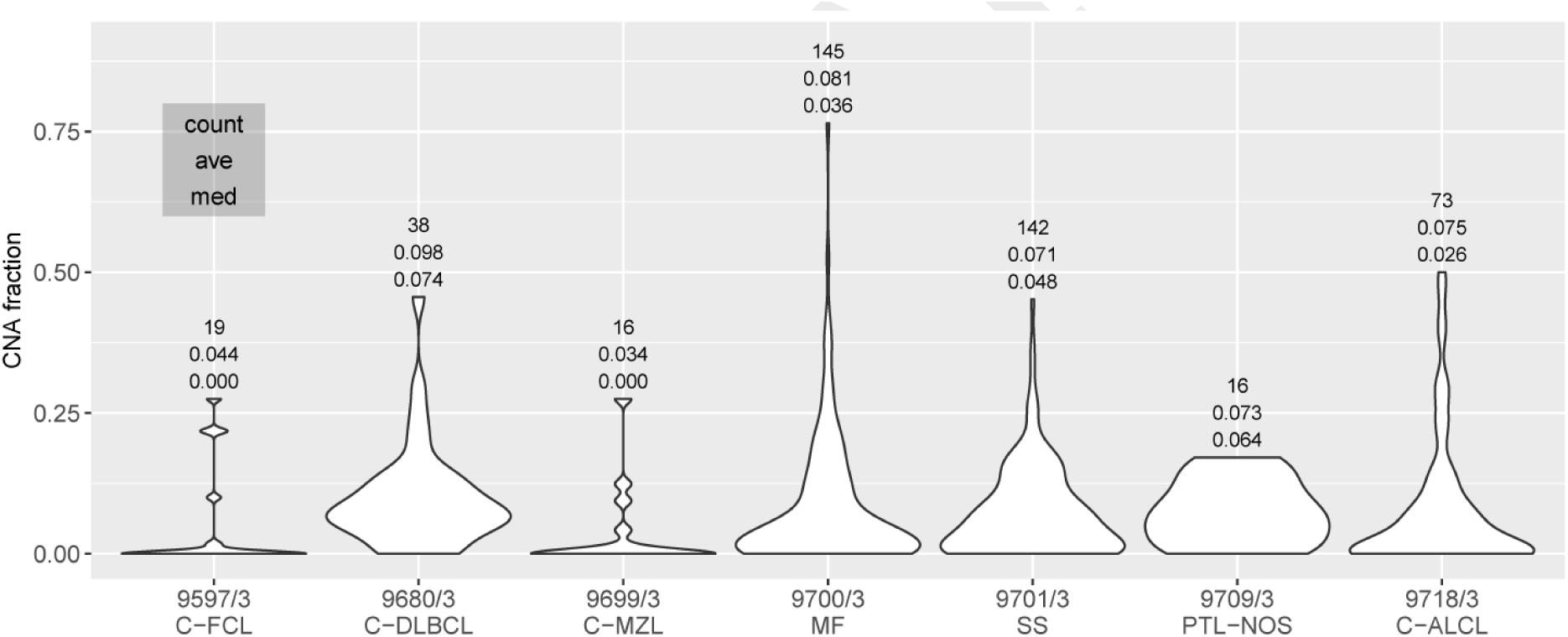
Fraction of genomic imbalances per sample. Extent of CNA coverage in different CL entities, as fractional CNA genome coverage (i.e. the fraction of a sample’s genome showing a CNA). The violin plot depicts the sample specific distribution of CNA fractions in the respective entities. Numbers above the graphs represent (top down) 1. the number of patients in that category of CL; 2. the average / arithmetic mean of fractional CNA coverage and 3. the CNA fraction median.

The lowest overall amount of CNA was observed in indolent cutaneous B-NHL (cFCL &cMZL), while higher rates of genomic imbalances were observed in cTNHL and especially DLBCL, with an overall agreement between clinical aggressiveness of disease type and amount of genomic abnormalities.

We want to point out that in the future the analysis of distribution and the fraction of genomic imbalances can be a novel prognostic factor to predict the clinical progression of the lymphoma and potentially direct the therapeutic approach.

### M. Survival data and prognoses

In addition, we analyzed the effect of existing copy number aberration on disease progression and survival. For 86 patients (90 samples) with cutaneous lymphomas of different histologies (Table 4), both follow-up and survival data were available. All of them were profiled using cCGH. 15 patients were aged <50 years and 71 patients were aged >50 years.

**Table 4.**
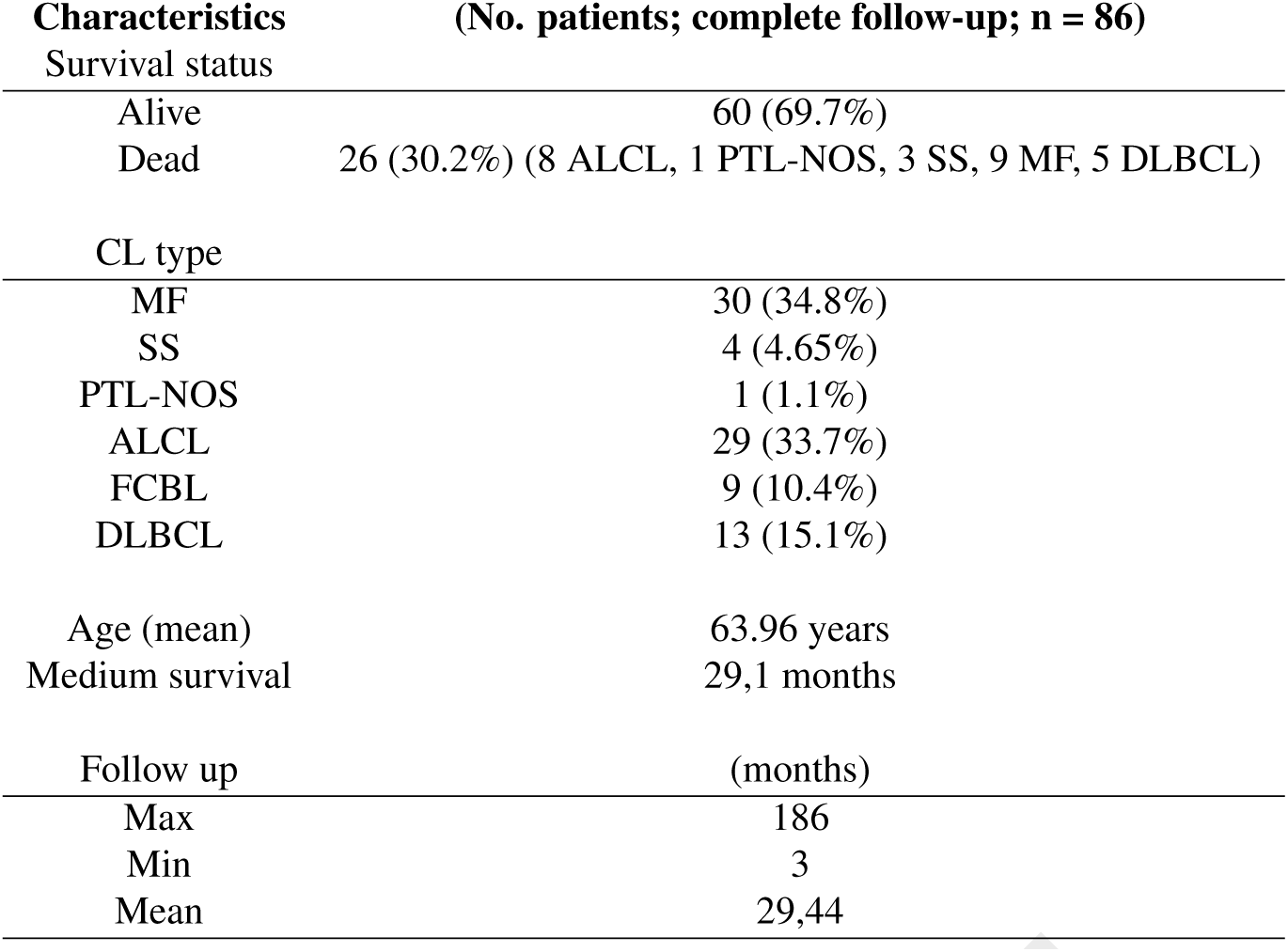
emographics and disease characteristics for 86 patients with CL

| Characteristics | (No. patients; complete follow-up; n = 86) |
| --- | --- |
| Survival status |  |
| Alive | 60 (69.7%) |
| Dead | 26 (30.2%) (8 ALCL, 1 PTL-NOS, 3 SS, 9 MF, 5 DLBCL) |
| CL type |  |
| MF | 30 (34.8%) |
| SS | 4 (4.65%) |
| PTL-NOS | 1 (1.1%) |
| ALCL | 29 (33.7%) |
| FCBL | 9 (10.4%) |
| DLBCL | 13 (15.1%) |
| Age (mean) | 63.96 years |
| Medium survival | 29,1 months |
| Follow up | (months) |
| Max | 186 |
| Min | 3 |
| Mean | 29,44 |

In a study of 20 cases of SS Vermeer *et al.* reported a median survival time of 31 months (17). In addition to this, some authors demonstrated that a higher number of CNAs resulted in a shorter overall survival (33, 34).

As shown in Table 5, MF has the largest number of CNAs. The most affected chromosomes were 6, 7 and 17. On chromosome 6 the most of CNAs are found in ALCL; whereas on chromosome 7 and 17, high number of CNAs are developed in MF.

**Table 5.** Number of CNAs per chromosome per disease in 86 patients (90 samples)

| CHR | ALCL | PTL-NOS | DLBCL | FCBL | MF | SS | Total |
| --- | --- | --- | --- | --- | --- | --- | --- |
| 1 | 3 |  | 7 | 1 | 12 |  | 23 |
| 2 | 3 |  | 2 | 2 | 11 | 2 | 20 |
| 3 | 7 |  | 2 |  | 6 | 1 | 16 |
| 4 | 2 |  | 1 | 1 | 9 | 1 | 14 |
| 5 | 4 |  | 1 | 1 | 11 | 1 | 18 |
| 6 | 13 |  | 9 |  | 9 | 2 | 33 |
| 7 | 9 |  | 7 | 1 | 10 | 1 | 28 |
| 8 | 6 | 1 | 2 |  | 10 | 4 | 23 |
| 9 | 8 |  | 1 | 1 | 13 | 1 | 24 |
| 10 | 3 |  | 2 |  | 7 | 4 | 16 |
| 11 | 2 |  | 1 |  | 12 | 2 | 17 |
| 12 | 3 |  | 5 | 2 | 9 | 1 | 20 |
| 13 | 5 |  | 1 |  | 6 | 2 | 14 |
| 14 | 3 |  |  | 1 | 6 |  | 10 |
| 15 | 4 |  | 2 | 1 | 3 |  | 10 |
| 16 | 6 |  | 1 |  | 7 | 1 | 15 |
| 17 | 6 |  | 4 |  | 16 | 5 | 31 |
| 18 | 2 |  | 12 |  | 2 | 1 | 17 |
| 19 | 2 |  |  |  | 9 | 1 | 12 |
| 20 | 3 |  | 1 |  | 6 | 2 | 12 |
| 21 | 2 |  |  |  | 2 |  | 4 |
| 22 | 1 |  |  |  | 5 |  | 6 |
| X | 2 |  | 4 | 1 | 5 | 1 | 13 |
| Y | 1 |  |  |  | 4 |  | 5 |
| Total | 100 | 1 | 65 | 12 | 190 | 33 | 401 |

We analyzed CNAs in this lot of patients to compare the results of survival with the reports from literature. We had 90 samples (86 patients) (Figure 7) from which 60 samples (57 patients) had CNAs and 30 samples (29 patients) are negative for CNAs.

**Fig. 7.**
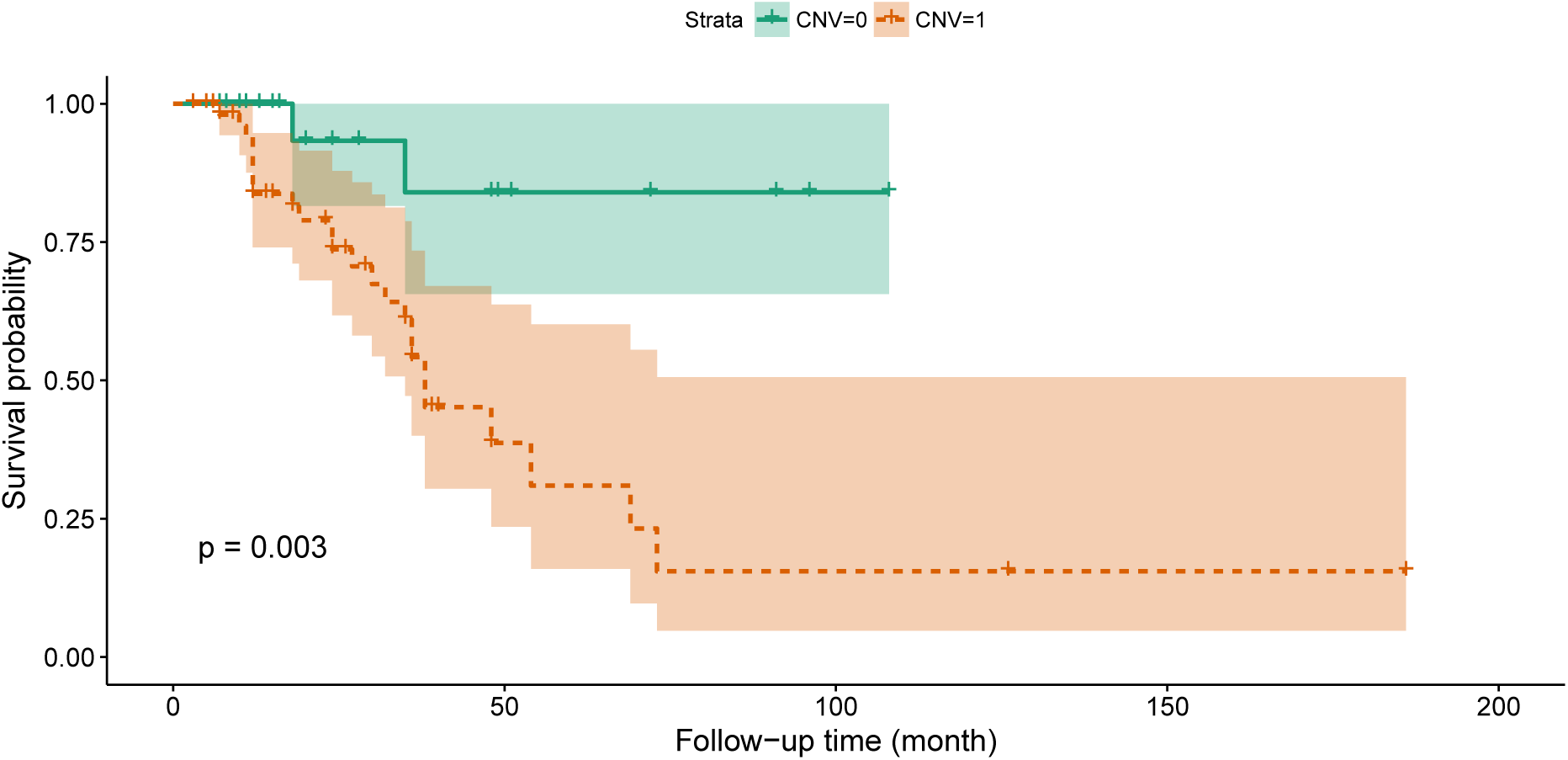
Kaplan-Meyer plot of patient survival with a maximum observation time of 186 months (86 patients of different histologies). The lower survival rate of patients with detected genomic instability reflects on the correlation of genome rearrangement events with more aggressive clinical course.

As we can see in the Table 6, the overall survival in patients without detected CNAs is superior to those with pronounced genomic instability. In the group with CNAs (57 patients), 24 died (42.1% with average follow up of 28.7 months). From the patients without CNAs (29 patients) two died (3.5%). We mention that from the one of the 2 CNA negative patients who died, a second sample showed a molecular progression with appearance of CNA 24 months after diagnosis. The survival curves obtained by using the Kaplan-Meyer method shown in Figure 7 represent the clinical course (maximum observation time 186 months) in those 86 patients. After 70 months, the survival rates seem to stabilize for both groups.

**Table 6.** CNA in 86 patients and overall survival

|  | CNA positive | CNA negative |
| --- | --- | --- |
| Alive | 33 | 27 |
| Dead | 24 | 2 |

Salgado *et al.* (33) in a multicenter study of 41 cases of tumor stage MF using a high resolution oligoarray comparative genomic hybridisation platform demonstrated that three specific chromosomal imbalances were associated with poor prognosis: gains of 8q24.21, deletion of 9p21.3 and 10q26qter.

We analyzed 30 MF cases metadata and we can observe gains of 8q24 in 6 cases but preponderant on 8q24.3, loses on 9p21 in 3 cases, and homogenous loses in 2 cases on 10q26. In the case of 9p21 and 10q26, all 5 patients died and in 8q24 gains, half of the patients survived.

Other factors that promoted a shorter survival rate were age (older than 60) and the presence of multiple cutaneous lesions (more than 2 sites) (33). In our case, out of 86 patients, 57 were equal or older than 60 and 23 (40.3%) died. Prochazkova M *et al.* in their study of common chromosomal imbalances in cutaneous CD30+ T cell lymphoma demonstrated that the number of chromosomal abnormalities was higher in the cases of patients with relapsing disease (mean number of changes 6.29). The samples obtained from these patients presented in most cases gains in chromosome 9 while chromosomes 6 and 18 were mostly affected by loss (6q and 18p) (35). Also, Fischer *et al.* in a study on 32 patients with CTCL showed that more aggressive tumors were presented with more pronounced chromosome imbalances (>/=5 CI meant a shorter survival). They also demonstrated that not all CNAs lead to a poorer prognosis, for example, the loss of 17p and the gain in chromosome 7 did not influence the prognosis. On the other hand, gains in 8q and loss of 6q and 13q were associated with a significantly shorter survival (34).

In our 86 patient analyses, 12 had deletions on 17p, out of which 6 had also gains on 17q and 1 deletion of 17q. From these 12 patients, 4 had MF, 3 SS, 3 ALCL and 2 DLBCL. Of these patients, 8 died (66%) with an average survival of 35.2 months. We observe that the medium survival in these cases is better than the one calculated for all 86 patients.

When it comes to gains in chromosome 7 and survival, we noticed that out of 86 patients 23 had gains in chromosome 7 Five were DLBCL, 7 were ALCL, 9 were MF, 1 SS and 1 FCBL. Eleven died (47%) with a medium survival of 24.8 months. We also looked over for gains in 8q to identify its influence on survival. Thirteen patients presented this CAN out of them 3 were ALCL, 2 DLBCL, 5 MF and 3 SS, of which 6 patients (46%) died with an average survival of 20.8 months. As shown, the average survival is lower than in the cases above but is not significantly reduced. Gains in 6q were encountered in 16 cases out of 86, 6 MF, 6 ALCL and 4 DLBCL. From these patients 10 died (62%) with a medium survival of 30.1 months. The survival in this case is greater than the mean for entire lot of patients (86) but the percentage of patients who did not survive versus those who did is significant. In the case of 13q loss, we analyzed 11 patients (4 MF, 5 ALCL, 1 DLBCL, 1 SS) out of 86, who had this CNA from which 7 died (63.6%) on 26.5 average month survival.

In Figure 8 we can easily notice the difference in survival rate between patients with or without CNV. The number of CNAs in patients who did not survive is much higher with frequent gains in 1, 6p, 7, 8q, 9qter, 14q, 15q, 16p, 17q, 19, 20q and 22q and frequent deletions on 3, 4q, 5q, 6q, 10q, 13, 17p.

**Fig. 8.**
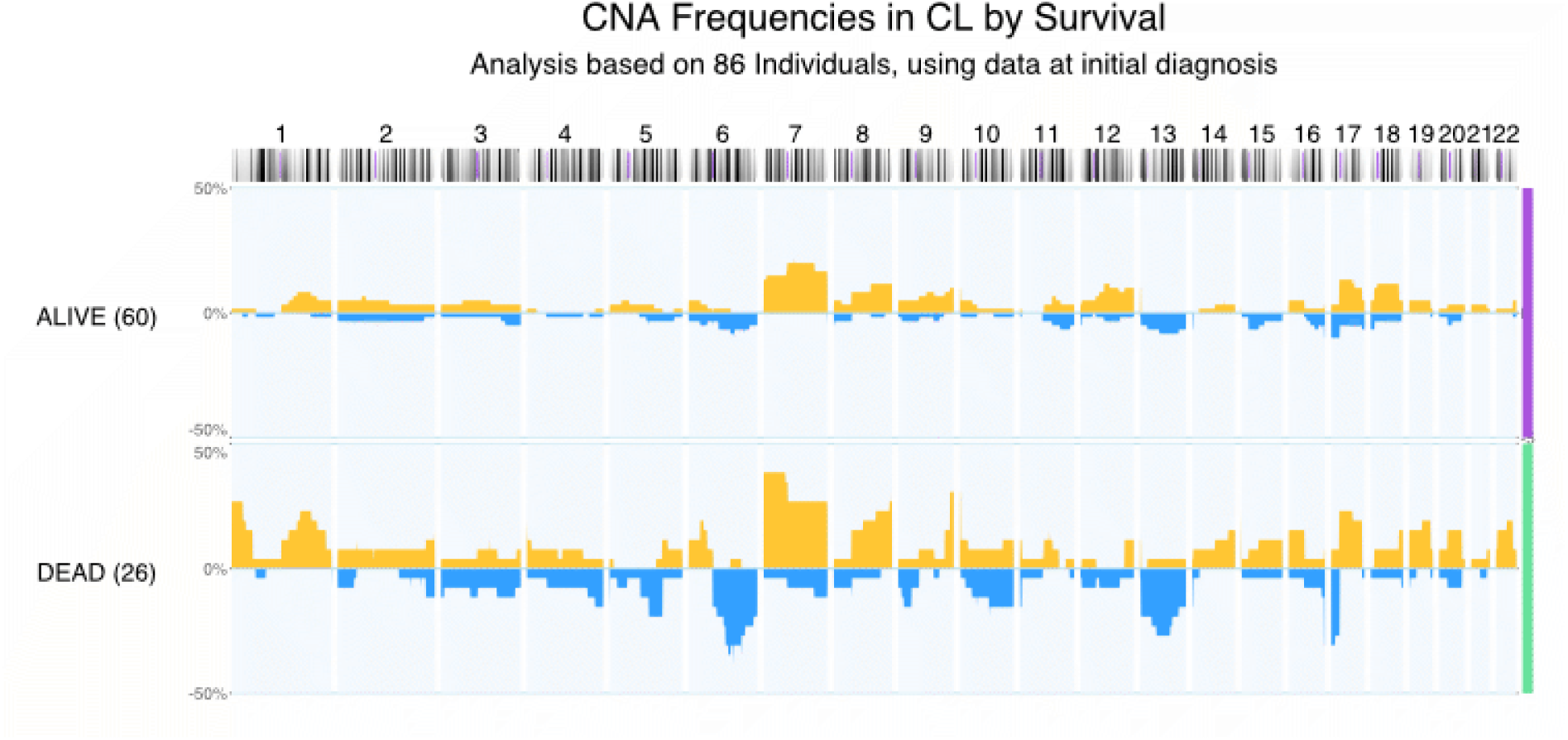
CNAs of 86 patients with different types of CL, separated for survival status.

On the other hand, the surviving group are presented with fewer CNAs. We report less frequent gains on 7, 8q, 9, 17q and 18.

We also correlate the average fraction of chromosomal imbalances with the survival. We observed more chromosomal imbalances in the patients who did not survive (average abfrac in deceased group 0.14 versus in surviving group 0.06).

## Discussion

Our data support the findings of individual genomic studies that the different types of CL are characterised by partially complex chromosomal imbalances, including structural and numerical abnormalities. While the small number of cases included in a single study frequently limits the comparative interpretation of molecular observations, our meta-analysis represents an approach to increase patient numbers for such comparisons and provides an integrative analysis of genomic imbalance patterns in CL.

Among the cases of MF the most frequent aberrations were represented by gains in chromosomes 7, 8, 17, 19, and 22. Previously, alterations of chromosomes 8 and 17 (detected by G-banding and FISH), had been associated with increased disease activity in MF (36). CGH had identified gains of chromosome 8 as a prognostic factor, together with deletions in chromosomes 9 and 10 (33). While those alterations could also be observed in SS, gains on chromosomes 19 and 22 were less common in SS patients, in contrast to copy number gains in chromosome 4 and deletions in chromosome 1, 10 (highly characteristics) and 22.

Studies have demonstrated that activation of PI3K/AKT signaling pathway due to PTEN alterations is rarely attributed to abnormalities in PTEN, PI3K and AKT1 genes, but still the presence of mutations in the PTEN gene (10q23.31) have a negative impact on the patient prognosis (37). Prominent among deregulated genes are those encoding MYC, MYC-regulating proteins, mediators of MYC-induced apoptosis, and IL-2 signaling pathway components (17).

In our meta-analysis, deletions on chromosome 10 were more frequent than gains on chromosome 12q, which had been reported previously in multiple studies among the most common cytogenetic aberration in different stages of CTCL, involving NAV 3 gene (POMFIL1) which may regulate IL-2 production of activated T-cells and may be implicated in cellular signaling and cell cycle regulation (38–40). Our study supports a possible role of chromosome 10 deletions in the pathogenesis of aggressive CTCL, especially SS, and encourages new studies to analyse the role of tumor suppressor genes found on this chromosome (e.g. MGMT and EBF3).

Duplications in chromosomes 8q (contains MYC) and 17q (contains STAT and ERBB3), proposed as early clonal events in tumor development (41) (17, 42), seems to be markers for CTCL, with very frequent alterations observed in MF and SS, but not in CBCL. ALCL presents with imbalances in all chromosomes, possibly reflecting a complex pathogenesis, but with most frequent CNA involving chromosome 7 (gains) as well as 6 and 13 (losses).

The most frequent abnormalities in CBCL are gains in chromosome 1, 2, 3, 7, 12, 13 and 18. DLBCL is characterized by a wide spectrum of imbalances, including a large number of deletions in chromosome 6 (also found in nodal lymphomas) (43, 44) as well as gains in chromosome 18 which contains the BCL2, a well-established anti-apoptotic effector involved in the pathogenesis of some nodal and extranodal lymphomas (45–49).

Limitations of the study include that the original data were produced over several years, with analyses performed on different platforms and in many different laboratories and mostly not being based on the “state-of-the-art” technologies (i.e. high-resolution genotyping arrays or whole-genome sequencing). Regarding the metadata, some of the studies were published before establishment of the two main classifications (WHO/EORTC and WHO); and the original diagnostic assignments didn’t follow their standardized criteria. Another important limitation is the small number of cases for some entities and therefore the risk of their biased representation. Also, this study was severely limited in correlating CNAs to clinical parameters such as “stage” due to the small number of cases with such information. Moreover, some rare types of CL could not be included in the analysis simply due to the absolute lack of genomic studies. However, we will provide support for future analyses by periodically updating the CL data accessible through the Progenetix and arrayMap repositories.

The results demonstrate that copy number gains and losses detected by genomic copy number profiling could be used at least as supporting information when classifying lymphomas into biologically and clinically distinct diseases or subtypes. Genomic copy number alterations have the potential to help diagnose or classify different disease entities, tumor subtypes, and even prognostic features.

In conclusion, CL are characterized by varying and sometimes complex CNA profiles, reflecting their variable pathogenesis. The identification of specific genomic imbalances could yield the critical insights of these intriguingly subjacent molecular events. Moreover, genomic signatures could support more accurate classifications, in an area which still represents a matter of debate.

## Acknowledgements

This study initiated from a project supported by the Swiss National Science Foundation (Grant/Award Number: IZERZ0_142305). We also thank UEFISCDI România for additional funding support.

## Abbreviations

ABFRAC: Average fractions of genomic imbalances per sample
aCGH: Array-based Comparative Genomic Hybridization
ALCL: anaplastic large cell lymphoma
B-NHL: B non-Hodgkin Lymphoma
CBCL: Cutaneous B cell Lymphomas
cCGH: Chromosomal Comparative Genomic Hybridization
CD4+ SMTL: Primary cutaneous CD4-positive small/medium T-cell lymphoma
CD8+ AECTCL: Primary cutaneous CD8-positive aggressive epidermotropic T-cell lymphoma
cFCL: Cutaneous Follicle centre lymphoma
CGD-TCL: Primary cutaneous gamma/delta T-cell lymphoma
CGH: Comparative Genomic Hybridization
CHR: Chromosome
CL: Cutaneous Lymphoma
cMZL: Cutaneous Marginal zone lymphoma
CNA: Copy number alterations
CNHL: Cutaneous non-Hodgkin Lymphoma
CNV: Copy number variations
CTCL: Cutaneous T cell Lymphomas
cTNHL: Cutaneous T non-Hodgkin Lymphoma
DLBCL: Diffuse large B-cell lymphoma
EORTC: European Organization for Research and Treatment of Cancer
FCBL: Primary cutaneous follicle centre lymphoma
HTLV-1: Human T-cell Lymphotropic Virus Type 1
LPP: Large Plaque Parapsoriasis
MALT lymphoma (MZBL): Extranodal marginal zone lymphoma of mucosa-associated lymphoid tissue
MF: Mycosis Fungoides
No: number
cALCL: Primary Cutaneous Anaplastic Large cell Lymphoma
PCL: Primary Cutaneous Lymphoma
PTL-NOS: Peripheral T-cell lymphoma not otherwise specified
SNP: Single Nucleotid Polymorphism
SS: Sézary Syndrome
TCRS: Primary cutaneous peripheral Tcell lymphoma
T-NHL: T non-Hodgkin Lymphoma
WHO: World Health Organisation

## Bibliography

1. R Willemze, H Kerl, W Sterry, E Berti, L Cerroni, S Chimenti, JL Diaz-Peréz, ML Geerts, M Goos, R Knobler, E Ralfkiaer, M Santucci, N Smith, J Wechsler, WA van Vloten, and CJ Meijer. Eortc classification for primary cutaneous lymphomas: a proposal from the cutaneous lymphoma study group of the european organization for research and treatment of cancer. Blood, 90(1):354–371., 1997.

2. R Willemze, ES Jaffe, G Burg, L Cerroni, E Berti, SH Swerdlow, E Ralfkiaer, S Chimenti, JL Diaz-Perez, LM Duncan, F Grange, NL Harris, W Kempf, H Kerl, M Kurrer, R Knobler, N Pimpinelli, C Sander, M Santucci, W Sterry, MH Vermeer, J Wechsler, S Whittaker, and CJ Meijer. Who-eortc classification for cutaneous lymphomas. Blood, 105(10):3768–3785., 2005.

3. E Campo, SH Swerdlow, NL Harris, S Pileri, H Stein, and ES Jaffe. The 2008 who classification of lymphoid neoplasms and beyond: evolving concepts and practical applications. Blood, 117(19):5019–5032., 2011.

4. SH Swerdlow, E Campo, SA Pileri, NL Harris, H Stein, R Siebert, R Advani, M Ghielmini, GA Salles, AD Zelenetz, and ES Jaffe. The 2016 revision of the world health organization classification of lymphoid neoplasms. Blood, 127(20):2375–2390., 2016.

5. HK Wong, A Mishra, T Hake, and P Porcu. Evolving insights in the pathogenesis and therapy of cutaneous t-cell lymphoma (mycosis fungoides and sezary syndrome). Br J Haematol, 155(2):150–166., 2011.

6. Health Quality Ontario. Extracorporeal photophoresis: an evidence-based analysis. Ontario health technology assessment series, 6:1, 2006.

7. B Vergier, A de Muret, M Beylot-Barry, L Vaillant, D Ekouevi, G Chene, A Carlotti, N Franck, P Dechelotte, P Souteyrand, P Courville, P Joly, M Delaunay, M Bagot, F Grange, S Fraitag, J Bosq, T Petrella, A Durlach, A De Mascarel, JP Merlio, and J Wechsler. Transformation of mycosis fungoides: clinicopathological and prognostic features of 45 cases. french study group of cutaneious lymphomas. Blood, 95(7):2212–2218., 2000.

8. DV Kazakov, G Burg, and W Kempf. Clinicopathological spectrum of mycosis fungoides. J Eur Acad Dermatol Venereol, 18(4):397–415., 2004.

9. M Setoyama, FA Kerdel, G Elgart, T Kanzaki, and JJ Byrnes. Detection of htlv-1 by polymerase chain reaction in situ hybridization in adult t-cell leukemia/lymphoma. Am J Pathol, 152(3):683–689., 1998.

10. G Burg, W Kempf, A Cozzio, J Feit, R Willemze, E S Jaffe, R Dummer, E Berti, L Cerroni, S Chimenti, JL Diaz-Perez, F Grange, NL Harris, DV Kazakov, H Kerl, M Kurrer, R Knobler, CJ Meijer, N Pimpinelli, E Ralfkiaer, R Russell-Jones, C Sander, M Santucci, W Sterry, SH Swerdlow, MH Vermeer, J Wechsler, and S Whittaker. Who/eortc classification of cutaneous lymphomas 2005: histological and molecular aspects. J Cutan Pathol, 32(10): 647–674., 2005.

11. JM McGregor, CC Yu, QL Lu, FE Cotter, DA Levison, and DM MacDonald. Posttransplant cutaneous lymphoma. J Am Acad Dermatol, 29(4):549–554., 1993.

12. M Nielsen, K Kaltoft, M Nordahl, C Röpke, C Geisler, T Mustelin, P Dobson, A Svejgaard, and N Odum. Constitutive activation of a slowly migrating isoform of stat3 in mycosis fungoides: tyrphostin ag490 inhibits stat3 activation and growth of mycosis fungoides tumor cell lines. Proc Natl Acad Sci U S A, 94(13):6764–6769., 1997.

13. D Jones, C O’Hara, MD Kraus, AR Perez-Atayde, A Shahsafaei, L Wu, and DM Dorfman. Expression pattern of t-cell-associated chemokine receptors and their chemokines correlates with specific subtypes of t-cell non-hodgkin lymphoma. Blood, 96(2):685–690., 2000.

14. JJ Scarisbrick, AJ Woolford, R Russell-Jones, and SJ Whittaker. Allelotyping in mycosis fungoides and sézary syndrome: common regions of allelic loss identified on 9p, 10q, and 17p. J Invest Dermatol, 117(3):663–670., 2001.

15. KD Wu and ER Hansen. Shortened telomere length is demonstrated in t-cell subsets together with a pronounced increased telomerase activity in cd4 positive t cells from blood of patients with mycosis fungoides and parapsoriasis. Exp Dermatol, 10(5):329–336., 2001.

16. L Tracey, R Villuendas, AM Dotor, I Spiteri, P Ortiz, JF Garcia, JL Peralto, M Lawler, and MA Piris. Mycosis fungoides shows concurrent deregulation of multiple genes involved in the tnf signaling pathway: an expression profile study. Blood, 102(3):1042–1050., 2003.

17. MH Vermeer, R van Doorn, R Dijkman, X Mao, S Whittaker, PC van Voorst Vader, MJ Gerritsen, ML Geerts, S Gellrich, O Söderberg, KJ Leuchowius, U Landegren, JJ Out-Luiting, J Knijnenburg, M Ijszenga, K Szuhai, R Willemze, and CP Tensen. Novel and highly recurrent chromosomal alterations in sézary syndrome. Cancer Res, 68(8):2689–2698., 2008.

18. L van der Fits, MS van Kester, Y Qin, JJ Out-Luiting, F Smit, WH Zoutman, R Willemze, CP Tensen, and MH Vermeer. Microrna-21 expression in cd4+ t cells is regulated by stat3 and is pathologically involved in sézary syndrome. J Invest Dermatol, 131(3):762–768., 2011.

19. YC Tang and A Amon. Gene copy-number alterations: a cost-benefit analysis. Cell, 152(3): 394–405., 2013.

20. DF Stroup, JA Berlin, SC Morton, I Olkin, GD Williamson, D Rennie, D Moher, BJ Becker, TA Sipe, and SB Thacker. Meta-analysis of observational studies in epidemiology: a proposal for reporting. meta-analysis of observational studies in epidemiology (moose) group. JAMA, 283(15):2008–2012., 2000.

21. L Karenko, S Sarna, M Kähkönen, and A Ranki. Chromosomal abnormalities in relation to clinical disease in patients with cutaneous t-cell lymphoma: a 5-year follow-up study. Br J Dermatol, 148(1):55–64., 2003.

22. M Prochazkova, E Chevret, G Mainhaguiet, J Sobotka, B Vergier, MA Belaud-Rotureau, M Beylot-Barry, and JP Merlio. Common chromosomal abnormalities in mycosis fungoides transformation. Genes Chromosomes Cancer, 46(9):828–838., 2007.

23. Genetic home references. https://ghr.nlm.nih.gov/gene/MYC. Accessed: 2018-06-21.

24. S Geller, TN Canavan, M Pulitzer, AJ Moskowitz, and PL Myskowski. Alk-positive primary cutaneous anaplastic large cell lymphoma: a case report and review of the literature. Int J Dermatol, 57(5):515–520., 2018.

25. Y Zeng and AL Feldman. Genetics of anaplastic large cell lymphoma. Leuk Lymphoma, 57 (1):21–27., 2016.

26. Madhu P Menon, Stefania Pittaluga, and Elaine S Jaffe. The histological and biological spectrum of diffuse large b-cell lymphoma in the who classification. Cancer journal (Sudbury, Mass.), 18(5):411, 2012.

27. M Covington, D Cassarino, and F Abdulla. Alk expression is a rare finding in mycosis fungoides. Am J Dermatopathol, 39(5):342–343., 2017.

28. U Wehkamp, I Oschlies, I Nagel, J Brasch, M Kneba, A Günther, W Klapper, and M We-ichenthal. Alk-positive primary cutaneous t-cell-lymphoma (ctcl) with unusual clinical presentation and aggressive course. J Cutan Pathol, 42(11):870–877., 2015.

29. X Mao, D Lillington, JJ Scarisbrick, T Mitchell, B Czepulkowski, R Russell-Jones, B Young, and SJ Whittaker. Molecular cytogenetic analysis of cutaneous t-cell lymphomas: identification of common genetic alterations in sézary syndrome and mycosis fungoides. Br J Dermatol, 147(3):464–475., 2002.

30. R van Doorn, MS van Kester, R Dijkman, MH Vermeer, AA Mulder, K Szuhai, J Knijnenburg, JM Boer, R Willemze, and CP Tensen. Oncogenomic analysis of mycosis fungoides reveals major differences with sezary syndrome. Blood, 113(1):127–136., 2009.

31. F Gallardo, B Bellosillo, B Espinet, RM Pujol, T Estrach, O Servitje, V Romagosa, C Barranco, S Boluda, M García, F Solé, A Ariza, and S Serrano. Aberrant nuclear bcl10 expression and lack of t(11;18)(q21;q21) in primary cutaneous marginal zone b-cell lymphoma. Hum Pathol, 37(7):867–873., 2006.

32. M Lima. Cutaneous primary b-cell lymphomas: from diagnosis to treatment. An Bras Dermatol, 90(5):687–706., 2015.

33. R Salgado, O Servitje, F Gallardo, MH Vermeer, PL Ortiz-Romero, MB Karpova, MC Zipser, C Muniesa, MP García-Muret, T Estrach, M Salido, J Sánchez-Schmidt, M Herrera, V Romagosa, J Suela, BI Ferreira, JC Cigudosa, C Barranco, S Serrano, R Dummer, CP Tensen, F Solé, RM Pujol, and B Espinet. Oligonucleotide array-cgh identifies genomic subgroups and prognostic markers for tumor stage mycosis fungoides. J Invest Dermatol, 130(4): 1126–1135., 2010.

34. TC Fischer, S Gellrich, JM Muche, T Sherev, H Audring, H Neitzel, P Walden, W Sterry, and H Tönnies. Genomic aberrations and survival in cutaneous t cell lymphomas. J Invest Dermatol, 122(3):579–586., 2004.

35. M Prochazkova, E Chevret, M Beylot-Barry, J Sobotka, B Vergier, M Delaunay, M Turmo, J Ferrer, P Kuglik, and JP Merlio. Chromosomal imbalances: a hallmark of tumour relapse in primary cutaneous cd30+ t-cell lymphoma. J Pathol, 201(3):421–429., 2003.

36. L Karenko, E Hyytinen, S Sarna, and A Ranki. Chromosomal abnormalities in cutaneous t-cell lymphoma and in its premalignant conditions as detected by g-banding and interphase cytogenetic methods. J Invest Dermatol, 108(1):22–29., 1997.

37. E Papadavid, P Korkolopoulou, G Levidou, AA Saetta, T Papadaki, M Siakantaris, V Nikolaou, A Oikonomidi, I Chatziandreou, L Marinos, A Kolialexi, A Stratigos, D Rigopoulos, A Psyrri, E Patsouris, and C Antoniou. In situ assessment of pi3k and pten alterations in mycosis fungoides: correlation with clinicopathological features. Exp Dermatol, 23(12): 931–933., 2014.

38. L Karenko, S Hahtola, S Päivinen, R Karhu, S Syrjä, M Kähkönen, B Nedoszytko, S Kytölä, Y Zhou, V Blazevic, M Pesonen, H Nevala, N Nupponen, H Sihto, I Krebs, A Poustka, J Roszkiewicz, K Saksela, P Peterson, T Visakorpi, and A Ranki. Primary cutaneous t-cell lymphomas show a deletion or translocation affecting nav3, the human unc-53 homologue. Cancer Res, 65(18):8101–8110., 2005.

39. DA Batista, EC Vonderheid, A Hawkins, L Morsberger, P Long, KM Murphy, and CA Griffin. Multicolor fluorescence in situ hybridization (sky) in mycosis fungoides and sézary syndrome: search for recurrent chromosome abnormalities. Genes Chromosomes Cancer, 45 (4):383–391., 2006.

40. S Hahtola, S Tuomela, L Elo, T Häkkinen, L Karenko, B Nedoszytko, H Heikkilä, U Saarialho-Kere, J Roszkiewicz, T Aittokallio, R Lahesmaa, and A Ranki. Th1 response and cytotoxicity genes are down-regulated in cutaneous t-cell lymphoma. Clin Cancer Res, 12(16):4812–4821., 2006.

41. G Barba, C Matteucci, G Girolomoni, L Brandimarte, E Varasano, MF Martelli, and C Mecucci. Comparative genomic hybridization identifies 17q11.2 approximately q12 duplication as an early event in cutaneous t-cell lymphomas. Cancer Genet Cytogenet, 184(1): 48–51., 2008.

42. A Carbone, L Bernardini, F Valenzano, I Bottillo, C De Simone, R Capizzi, A Capalbo, F Romano, A Novelli, B Dallapiccola, and P Amerio. Array-based comparative genomic hybridization in early-stage mycosis fungoides: recurrent deletion of tumor suppressor genes bcl7a, smac/diablo, and rhof. Genes Chromosomes Cancer, 47(12):1067–1075., 2008.

43. Y Hayashi, SC Raimondi, AT Look, FG Behm, GR Kitchingman, C-H P4i, GK Rivera, and DL Williams. Abnormalities of the long arm of chromosome 6 in childhood acute lymphoblastic leukemia. Blood, 76(8):1626–1630., 1990.

44. D Millikin, E Meese, B Vogelstein, C Witkowski, and J Trent. Loss of heterozygosity for loci on the long arm of chromosome 6 in human malignant melanoma. Cancer Res, 51(20): 5449–5453., 1991.

45. M Bentz, CA Werner, H Döhner, S Joos, TF Barth, R Siebert, M Schröder, S Stilgenbauer, K Fischer, P Möller, and P Lichter. High incidence of chromosomal imbalances and gene amplifications in the classical follicular variant of follicle center lymphoma. Blood, 88(4): 1437–1444., 1996.

46. O Monni, H Joensuu, K Franssila, and S Knuutila. Dna copy number changes in diffuse large b-cell lymphoma–comparative genomic hybridization study. Blood, 87(12):5269–5278., 1996.

47. CA Werner, H Döhner, S Joos, LH Trümper, M Baudis, TF Barth, G Ott, P Möller, P Lichter, and M Bentz. High-level dna amplifications are common genetic aberrations in b-cell neoplasms. Am J Pathol, 151(2):335–342., 1997.

48. TF Barth, H Döhner, CA Werner, S Stilgenbauer, M Schlotter, M Pawlita, P Lichter, P Möller, and M Bentz. Characteristic pattern of chromosomal gains and losses in primary large b-cell lymphomas of the gastrointestinal tract. Blood, 91(11):4321–4330., 1998.

49. MC Simmonds, JP Higgins, LA Stewart, JF Tierney, MJ Clarke, and SG Thompson. Meta-analysis of individual patient data from randomized trials: a review of methods used in practice. Clin Trials, 2(3):209–217., 2005.

